# Postnatal interaction of size and shape in the human endocranium and brain structures

**DOI:** 10.1101/2025.01.20.633848

**Authors:** Kuranosuke Takagi, Osamu Kondo

## Abstract

The uniqueness of human brain growth and development has been considered promising to understand the advanced cognitive ability of humans. Compared with that of chimpanzees, the human endocranium undergoes several characteristic shape changes immediately after birth, which has been termed “endocranial globularization.” However, how the brain structures and surrounding neurocranium interact with each other during early development in the context of brain–neurocranium integration remains to be investigated. We observed shape changes in the human brain and endocranium during postnatal development using magnetic resonance imaging, and analyzed spatial constraints and interactions among subdivisions of the brain influencing endocranial morphology. Our results suggest that during postnatal development, the relative size changes of supratentorial and infratentorial regions and the cranial base largely constrain brain and endocranium shape. Specifically, a disproportionate increase in the size of the cerebellum relative to the cranial base affects infratentorial spatial packing constraints in neonates, causing inferoposterior expansion of the posterior cranial fossa and coronal reorientation of the petrous bone without lateral opening of the tentorium cerebelli. The dramatic size increase of the cerebellum relative to the cranial base immediately after birth is inferred to be characteristic of human development, which corresponds to previously observed bony shape changes.

## Introduction

The origin of the advanced cognitive ability of humans is an intriguing theme in anatomy and paleoanthropology. Comparative studies of the brains of hominoids, primates, and mammals have studied various features relatively extensively, such as brain mass relative to body mass (Jerison, 1977; Herculano-Houzel, 2007), the allometry of brain structures (Rilling & Insel, 1998, 1999; Barton & Harvey, 2000; Rilling, 2006), cortical connectivity (Rilling et al., 2008), and shape (Aldridge, 2011; Gómez-Robles et al., 2014, Bruner et al., 2017). Scholars have fiercely debated the uniqueness of the modern human brain. Some studies have focused on finding human-specific structures in the frontal lobe (Semendeferi et al., 2002; Schoenemann et al., 2005; Barton & Venditti, 2013; Passingham & Smaers, 2014), whereas others have argued that the human cognitive advantage stems from the large size of the human brain, which is a scaled-up version of the primate brain, with the latter already being economical in terms of neuronal density (Herculano-Houzel, 2009, 2012).

In paleoneurology, where the evolution of the hominin brain has been discussed on the basis of the fossil evidence, the endocast morphology of the hominin crania has been analyzed because the brain itself does not remain in fossil specimens. Paleoneurologists have extensively studied the uniqueness of the modern human endocast in terms of various features, such as cranial capacity (Godfrey & Jacobs, 1981), brain mass relative to body mass (Ruff et al., 1997), sulcul patterns (Holloway, 1983; Falk, 1985), endocast shape (Bruner et al., 2003; Bastir et al., 2011) and the estimated volumes of brain regions (Weaver, 2005; Kochiyama et al., 2018). In addition, developmental patterns of the cranial base and the endocranial morphology have been compared between humans and great apes (Dean & Wood, 1984; Lieberman & McCarthy, 1999; Neubauer et al., 2010; Zollikofer et al., 2017). However, our understanding of human brain evolution is still limited because endocast morphology does not fully correspond to the internal brain structure (Bruner et al., 2015; Alatorre Warren et al, 2019; Dumoncel et al., 2021), and the evolutionary and developmental mechanisms linking the brain and the braincase have not been clarified.

A series of hypotheses have been proposed concerning the evolution and development of the brain and the braincase. Following conjecture concerning the role of surrounding organs as functional matrices in neurocranial development (Moss and Young, 1960), the “spatial packing” hypothesis of neurocranial integration has recently been supported. The spatial packing hypothesis posits that the spatial constraints caused by the relative sizes of the neurocranium, the cranial base, and the face influence their morphologies (Gould, 1977; Ross & Ravosa, 1993; Zollikofer et al., 2017). Gould (1977) hypothesized that the spatial constraint imposed by the tradeoff between increased brain size and a relatively shortened cranial base causes cranial base flexion. Interspecific variation among primates supports this hypothesis, where species with relatively large brains have a more flexed cranial base (Ross & Ravosa, 1993). In terms of ontogeny, similar endocranial shape changes have been observed among great apes and recent and fossilized *Homo sapiens*, with enlargement of the cranial base and face region relative to the endocast during late ontogeny (Zollikofer et al., 2017, 2022), where the endocranial shape difference between humans and great apes also relates to this relative size change. Because similar trends were observed in humans and great apes, it was argued that the observed spatial packing constraints can be considered as a common feature of neurocranial integration in both ontogenetic and evolutionary terms (Zollikofer et al., 2017). To assess this hypothesis, how neurocranial shape variation is related to the degree of encephalization has been investigated (Ross & Ravosa, 1993; Jeffery & Spoor, 2002; Jeffery, 2003; Zollikofer et al., 2017, 2022; Jeffery et al., 2022). Relative encephalization was calculated as the ratio of the cube root of endocranial volume divided by the sagittal cranial base length (Ross & Ravosa, 1993; Jeffery & Spoor, 2002; Jeffery, 2003), the cube root of endocranial volume divided by the basicranial centroid size (CS) (Jeffery et al., 2022), or the CS of the endocranium divided by that of the cranial base and face (Zollikofer et al., 2017, 2022). By contrast, ontogenetic changes in the neurocranium in humans and a few primate species are reportedly inconsistent with the spatial packing hypothesis. Prenatally, the cranial base retroflexes in humans, macaques, and howler monkeys, although it should be flexed under the general spatial packing hypothesis as the endocranial volume becomes larger relative to the cranial base length (Jeffery & Spoor, 2002; Jeffery, 2003). During the early postnatal period in humans, the cranial base flexes rapidly, and the petrous bone’s orientation changes coronally (Dean & Wood, 1984; Lieberman & McCarthy, 1999), with little change in the relative encephalization index (Zollikofer et al., 2017). Because this is not observed in great apes, it is assumed to be a human-specific developmental feature (Dean & Wood, 1984; Lieberman & McCarthy, 1999; Zollikofer et al., 2017). Furthermore, Neubauer et al. (2010) observed expansion of the parietal and cerebellar regions in human neonates, but not in chimpanzees, during postnatal development, with this process being referred to as “globularization.” More recently, not only humans but also gorillas and orangutans were found to commonly exhibit relative expansion of the posterior cranial fossa, the temporal pole, and the orbitofrontal region during early postnatal development, whereas chimpanzees and bonobos did not (Zollikofer et al., 2017). The presence of neurocranial globularization in Neanderthals has been debated (Gunz et al., 2010, 2012; Ponce de León et al., 2016). Although it is assumed to be related to the development of the brain and brain structures given that the brain grows rapidly at the same time (Neubauer et al., 2010), the mechanisms of this shape change are not fully understood.

Within the endocast, the brain is compartmentalized and supported by the dura mater, including the falx cerebri, the falx cerebelli, and the tentorium cerebelli (Moss, 1958; Moss & Young, 1960, Standring et al., 2005). The falx cerebri separates the two cerebral hemispheres along the great longitudinal fissure in the sagittal plane and is attached to the cranial base anteriorly, and to the internal occipital protuberance posteriorly (Standring et al., 2005). The tentorium cerebelli divides the endocranial cavity into supratentorial and infratentorial regions, which contain the forebrain and hindbrain, respectively (Standring et al., 2005). In addition, the falx cerebri and the tentorium cerebelli reportedly have an essential role in supporting the heavy supratentorial cerebral hemispheres against gravity (Bull, 1969). These meningeal structures determine the direction of brain expansion during development and affect the endocranial shape (Moss & Young 1960; Friede, 1981), where compartmentalized brain regions may grow disproportionately.

As for the growth of the cerebellum, it expands early in the first and second years after birth (Knickmeyer et al., 2008). The relative expansion of the cerebellum may be associated with the expansion of the posterior cranial fossa observed in globularization (Neubauer et al., 2009), although this developmental pattern of the brain seems to be preserved across primate species; macaques also exhibit relative expansion of the cerebellum after birth (Scott et al., 2016). Overall, the dura mater is assumed to limit endocranial shape changes during the course of brain regional expansion before fusion of the neurocranium.

In terms of disproportionate growth where the dural structure acts as a border, it is worth noting the “infratentorial spatial packing” problem, where it is hypothesized that the relatively short posterior cranial base limits sagittal expansion of the cerebellum under the tentorium cerebelli such that it expands laterally and causes a coronal orientation of the petrosal bone in humans (Dean, 1988; White, 2005). Jeffery and Spoor (2002) tested this hypothesis in prenatal human fetuses, reporting no significant correlations between the relative volume of the infratentorial space and changes in cranial base flexion or petrosal bone orientation. Previous studies of human fetuses showed that the tentorium cerebelli changes its orientation in the sagittal plane with the relative expansion of the cerebral volume during the same period (Moss et al., 1956; Jeffery, 2002; Matsunari et al., 2023), and this structural change potentially influences the degree of the infratentorial spatial constraint during prenatal development. In adult primate species, Ross & Ravosa (1993) showed a non-significant relationship between the absolute cerebellar size and cranial base flexion.

To fill the knowledge gap and understand the developmental relationship between the neurocranium and the brain, we need to understand the growth patterns of the brain, including of internal brain structures such as the dura mater, during early development. However, although most previous studies have analyzed the developmental patterns of bony shape (Dean & Wood, 1984; Lieberman & McCarthy, 1999; Neubauer et al., 2010; Zollikofer et al., 2017), few studies have focused on the growth patterns of soft tissue structures inside the endocast or on the interactions between bone and soft tissue structures at that time (e.g., Rehder et al., 2016; Jeffery et al., 2022). The aim of this study is to observe developmental patterns of the brain, including the dura mater and endocranial structures, during the postnatal ontogeny in humans and to elucidate structural relationships related to previously reported bony shape changes, such as globularization. We observed magnetic resonance images of humans, covering the period from shortly after birth to adulthood, and characterized the morphologies of soft tissue structures, the cranial base, and the endocast via landmark-based geometric morphometrics. Additionally, we measured the sizes of the cranial base and the supra- and infratentorial spaces, as well as the angle of the cranial base flexion, the petrous orientation, and the dura mater configuration. The results shed light on how regional spatial constraints, such as infratentorial spatial packing, influence bony shapes and phenomena like globularization during the early postnatal period.

## Methods

### Data preparation

Modern human samples were obtained from the two open-access archives of the National Institute of Mental Health (NIMH) (Table 1), namely a battery of neurocranial magnetic resonance images of healthy Americans, from birth to adulthood, including 12 longitudinal specimens (25 images) and 77 cross-sectional specimens. We chose datasets to which a filter was applied to correct spatial distortion caused by the nonlinearity of the gradient field. In addition, we excluded facial morphology from the target cranial region because the images in one of the datasets (NIMH data archive ID: 1151) were blurred around the facial region. Concerning age categories, ontogenetic age groups were distinguished on the basis of chronological age, corresponding to the maxillary dental eruption stage (Sandler, 1944; AlQahtani et al., 2010), to allow comparison with previous ontogenetic studies (Table 2): up to 3 weeks after birth (stage 1), 3 weeks to 0.8 years (9.6 months) after birth (stage 2), incomplete deciduous dentition (stage 3), complete deciduous dentition (stage 4), and first/second permanent molar fully erupted (stages 5/6). In several specimens with multiple longitudinal images, one image was selected for each stage/age group, such that the images were treated in the same way as the cross-sectional ones.

**Table 1.** The MRI data repositories used in this study.

| Name | Age range | *N*<br>sampled | MR<br>sequence | Voxel resolution<br>(mm) | references |
| --- | --- | --- | --- | --- | --- |
| NIMH data<br>archive<br>(ID:1151) | 0-20y | 57 | T1 | 0.9375-1(x,y),<br>0.9375-3(z) | (Evans & Brain<br>Development<br>Cooperative<br>Group, 2006) |
| NIMH data<br>archive<br>(ID:2607) | 2-22y | 45 | T1 | 0.9375-1(x,y,z) | (Jernigan et al.,<br>2016) |

**Table 2.** The analyzed samples by age stage and sex.

| Stage | Age range | Dental stage | N total<br>(error) | N_unknown | N_male | N_female |
| --- | --- | --- | --- | --- | --- | --- |
| 1 | 0-3 weeks | No teeth erupted | 5 (3) | 5 (3) |  |  |
| 2 | 3 weeks-0.8 years |  | 13 (3) | 13 (3) |  |  |
| 3 | 0.8-2.5 years | Incomplete<br>deciduous dentition | 19 (3) | 19 (3) |  |  |
| 4 | 2.5-6.5 years | complete deciduous<br>dentition | 28 (3) | 8 (2) | 10 (1) | 10 |
| 5 | 6.5-13.5 years | M1 fully erupted | 17 (3) | 6 (1) | 2 (1) | 9 (1) |
| 6 | >= 13.5 years | M2 fully erupted | 20 (3) | 6 (3) | 6 | 8 |
| total |  |  | 102 (18) | 57 (15) | 18 (2) | 27 (1) |

Virtual endocasts were extracted for most of the specimens using the segmentation protocol in the SPM software package (Ashburner and Friston, 2005), where T1-weighted images were segmented into gray matter, white matter, and cerebrospinal fluid regions. For young specimens (stages 1–3), to which the same segmentation protocol could not be applied, the skull and outer soft tissue were stripped using the Brain Extraction Tool of Smith (2002). This procedure was performed because the human neonate and adult brain differ in tissue contrast because of the dominance of nonmyelinated white matter in neonates (Barkovich et al., 1988; Bird et al., 1989; Despotović et al., 2015). The extracted virtual endocasts were manually corrected by comparing them with magnetic resonance images in ITK-SNAP (www.itksnap.org; Yushkevich et al., 2006). Then, three-dimensional (3D) surface models were generated in Analyze 14.0 (Mayo Clinic, Biomedical Imaging Resource Core, Rochester, MN, USA), with surface smoothing performed in Geomagic Wrap (3D Systems, Inc., Rock Hill, SC, USA)

Landmarks were digitized on aligned T1-weighted images, and on the surface of the extracted 3D endocasts. The two sets of landmarks of different modalities were then merged, with the same anatomical landmarks used as a guide for each specimen. T1-weighted images were aligned on the basis of anatomy, with the transverse plane being parallel to the anterior commissure–posterior commissure line and the sagittal plane being parallel to the interhemispheric fissure. We digitized 37 anatomical landmarks both inside and outside the brain using Analyze 10.0 and Stratovan Checkpoint (Stratovan Corporation, Sacramento, CA, USA) (Table 3, Figure 1). In addition, we included 136 sliding semilandmarks (SLMs) in the 3D surface model, including 20 curve, 98 cerebral, and 18 cerebellar surface SLMs (Table 3, Figures 1 and 2(a)). Curve SLMs were captured between anatomical landmarks in each specimen, and surface SLMs were defined using one specimen as a template. After combining the two sets of landmarks, they were analyzed using the Morpho R package (Schlager 2017). The curve SLMs were resampled equidistantly between the fixed anatomical landmarks (Botton-Divet et al., 2016). The surface SLMs were warped from the template to each specimen by deforming the template onto a target specimen using the thin-plate spline algorithm, on the basis of the anatomical landmarks and the curve SLMs, with the warped surface SLMs then being projected onto the target surface. To establish geometric homology in the positions of the SLMs by minimizing bending energy, the curve and surface SLMs were slid along the tangents to the curves and the tangent planes to the surface, respectively (Gunz et al., 2005). Because we intended to capture 3D variation as a symmetric object, we divided it into symmetric and asymmetric components using MorphoJ (Klingenberg et al., 2002) and used only the symmetric component. For the symmetric component, we performed generalized Procrustes analysis to superimpose the landmarks and then subjected the variation relative to the average landmarks to principal component analysis (PCA) using MorphoJ.

**Figure 1.**
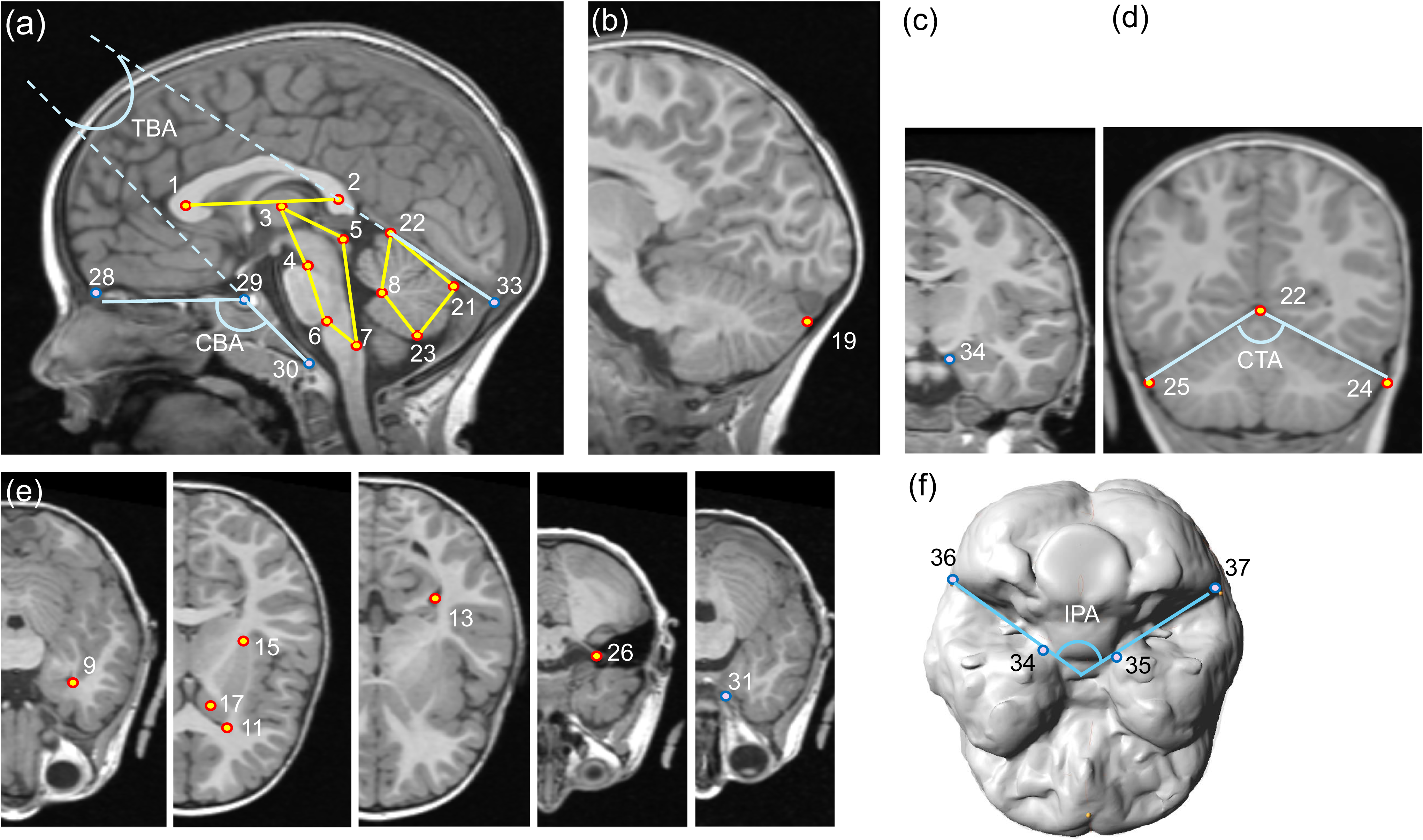
Landmarks and wireframe diagram of the angle measurements. (a) Midsagittal image; (b) sagittal image; (c) coronal image; (d) coronal tentorial angle (CTA); (e) transversal images (half); (f) inferior surface of the virtual endocast.

**Figure 2.**
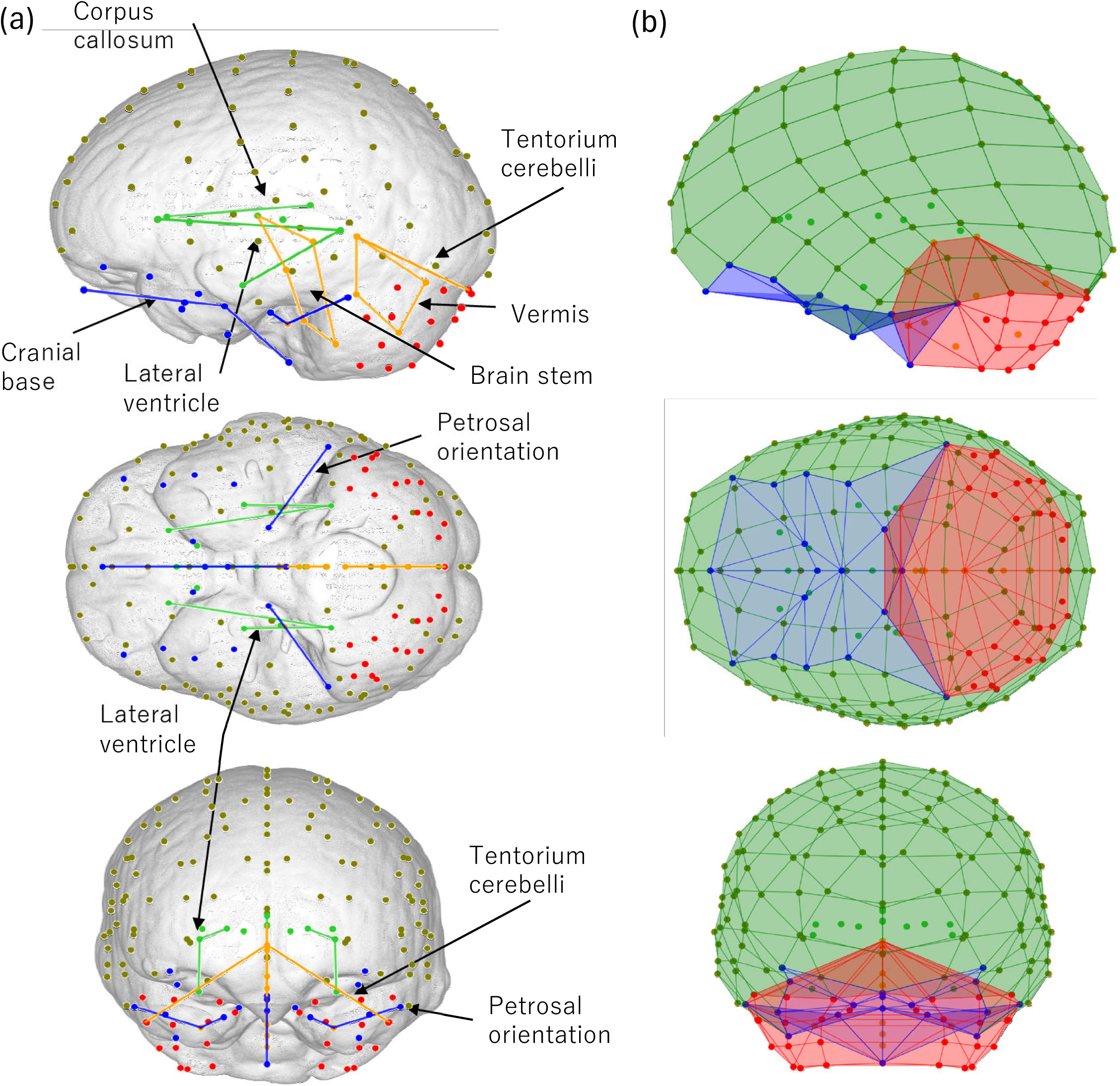
Landmark set and division of surface and volume mesh. (a) Landmarks and semilandmarks on the virtual endocast surface; (b) surface area and volume boundaries used to calculate the relative sizes. Colors indicate boundaries: cranial base surface area (region A, blue), infratentorial volume (region B, red), and supratentorial volume (region C, green).

**Table 3.** Landmarks and measurements.

|  | Landmark no. | Name | definition | Region | Landmark type | No. per set |
| --- | --- | --- | --- | --- | --- | --- |
| Brain landmarks | 1 | genu | The center of the genu of the corpus callosum | C | Fixed LM | 1 |
|  | 2 | splenium | The center of the splenium of the corpus callosum | C | Fixed LM | 1 |
|  | 3 | TH | The center of the thalamus | C | Fixed LM | 1 |
|  | 4 | IRI | The most posterior point of the interpeduncular fossa | B | Fixed LM | 1 |
|  | 5 | IC | The center of the inferior colliculus | B | Fixed LM | 1 |
|  | 6 | FC | The most posterior point of the foramen cecum of the brainstem | B | Fixed LM | 1 |
|  | 7 | CS | The pit point of the calamus scriptorius | B | Fixed LM | 1 |
|  | 8 | Forth ventricle | The apex of the median dorsal recess of the fourth ventricle | B | Fixed LM | 1 |
|  | 9,10 | Lateral ventricle inferior horn | The most anterior point of the inferior horn of the lateral ventricle | C | Fixed LM | 2 x 1 |
|  | 11,12 | Lateral ventricle anterior horn | The most anterior point of the anterior horn of the lateral ventricle | C | Fixed LM | 2 x 1 |
|  | 13,14 | Lateral ventricle trigone | The center of the trigone of the lateral ventricle | C | Fixed LM | 2 x 1 |
|  | 15,16 | Putamen | The most posterior point of the putamen | C | Fixed LM | 2 x 1 |
|  | 17,18 | Caudate nucleus head | The center of the head of the caudate nucleus | C | Fixed LM | 2 x 1 |
|  | 19,20 | Cerebellum posterior pole | The most posterior point of the cerebellum | B | Fixed LM | 2 x 1 |
|  | 21 | Most posterior point of cerebellar vermis | The most posterior point of the cerebellar vermis | B | Fixed LM | 1 |
|  | 22 | Most superior point of cerebellar vermis | The most superior point of the cerebellar vermis | B | Fixed LM | 1 |
|  | 23 | Most inferior point of cerebellar vermis | The most inferior point of the cerebellar vermis | B | Fixed LM | 1 |
|  | 24,25 | Cerebellar most lateral point | The most lateral point of the cerebellum | B | Fixed LM | 2 x 1 |
|  | 26,27 | Internal acoustic meatus | The point on the nerve at the internal acoustic meatus | B | Fixed LM | 2 x 1 |
| Cranial base landmarks | 28 | CG | The most anteroinferior point of the brain or the dura mater at the clista galli | A | Fixed LM | 1 |
|  | 29 | sella | The centor of the pituitary gland | A | Fixed LM | 1 |
|  | 30 | basion | The most anterior point of the foramen magnum | A | Fixed LM | 1 |
|  | 31,32 | optic canal | The point on the nerve at the optic canal | A | Fixed LM | 2 x 1 |
|  | 33 | IOP | The most anterior point of the internal occipital protuberance | A | Fixed LM | 1 |
|  | 34,35 | Petrosal apex | The apex of the petrous bone | A | Fixed LM | 2 x 1 |
|  | 36,37 |  | The point at the junction of the transverse sinus and the petrous pyramid | A | Fixed LM | 2 x 1 |
| Endocast landmarks |  | Transverse sinus | The curve semilandmarks between LM 33 and LM 36 and between LM 33 and LM 37 | B | Curve SLM | 2 x 3 |
|  |  | Midsagittal cerebrum | The curve semilandmarks between LM 28 and LM 33 | C | Curve SLM | 14 |
|  |  | Cerebral surface | The surface semilandmarks on the cerebral surface | A,C | Surface SLM | 2 x 49 |
|  |  | Cerebellar surface | The surface semilandmarks on the cerebellar surface | B | Surface SLM | 2 x 9 |
| Indices |  | Index of relative encephalization (IRE) | Cube root of the brain volume (B+C) divided by square root of the cranial base surface area (A). |  |  |  |
|  |  | Index of relative infratentorial encephalization (RIE) | Cube root of the infratentorial volume (B) divided by square root of the cranial base surface area (A). |  |  |  |
|  |  | Index of differential encephalization inside brain (IDE) | The infratentorial volume (B) divided by the supratentorial volume (C). |  |  |  |
|  |  | Cephalic index | The maximum width of the brain divided by the maximum length of the brain. |  |  |  |
| Angles |  | Cranial base angle (CBA) | Ventral angle between the anterior cranial base and the posterior cranial base (sella (No.29) -basion (No.30)). |  |  |  |
|  |  | Inter-petrosal angle (IPA) | Posterior angle between left and right line segments fitted through petrous bone apex (Nos.34, 35) and junction of transverse sinus and petrous pyramid (Nos.36, 37). |  |  |  |
|  |  | Tentorium-base angle (TBA) | Angle between the sagittal tentorium cerebelli orientation (Most superior point of cerebellar vermis (No.22) -IOP (No.33)) and the posterior cranial base (sella (No.29) -basion (No.30)). |  |  |  |
|  |  | Coronal tentorial angle (CTA) | Angle between the left and right line segments fitted through most superior point of cerebellar vermis (No.22) and left and right cerebellar most lateral points (Nos.24, 25) for each. |  |  |  |

### Relative sizes and angles inside the brain

To assess the previously proposed spatial constraint hypothesis, measures of relative size were calculated as the ratio of the sizes of regions of interest, including the cranial base (region A), the infratentorial region (region B), and the supratentorial region of the brain and the endocast (region C) (Table 3, Figure 2(b)). The size of the cranial base was given by the surface area of the 3D mesh created by the landmarks of the cranial base (Table 3, Figure 2(b), blue region). The infratentorial and supratentorial regions were quantified on the basis of volumes that encompassed the landmarks in these regions (Table 3, Figure 2(b); red and green regions, respectively). The surface area and volumes were calculated in MATLAB (MathWorks Inc., Natick, MA, USA). To test the general spatial packing hypothesis, the cube root of the whole brain volume (equal to the whole endocast; B + C) was divided by the square root of the cranial base (A) surface area as an index of relative encephalization (IRE). A very similar measure of relative encephalization has been used under the spatial packing hypothesis, where the global shape changes of the brain and the cranium are assumed to correlate with the IRE (Ross & Ravosa, 1993; Jeffery & Spoor, 2002; Jeffery, 2003; Zollikofer et al., 2017, 2022; Jeffery et al., 2022). The relative infratentorial encephalization (RIE) was calculated by dividing the cube root of the infratentorial (B) volume by the square root of the cranial base (A) surface area to test the infratentorial spatial packing hypothesis. Furthermore, the ratio of the infratentorial (B) volume and the supratentorial (C) volume was calculated as the index of differential encephalization inside the brain (IDE).

We also measured four angles and one index (Table 3) to observe the developmental patterns of the brain and cranial base shape. As a measure of cranial base flexion, the cranial base angle (CBA) was calculated between the anterior (sella [no. 29] to the most inferior point of the foramen caecum) and posterior cranial base (sella to basion [no. 30]). The definition of CBA in this study corresponds to measurement “CBA1” in Lieberman and McCarthy (1999), although a difference exists between the observations of soft and hard tissue. As a measure of the petrous bone’s orientation, the inter-petrosal angle (IPA) was calculated between the left and right line segments passing through the petrous bone apex (nos. 34 and 35) and the junction of the transverse sinus and petrous pyramid (nos. 36 and 37). As a measure of the sagittal orientation of the tentorium cerebelli relative to the posterior midsagittal cranial base, the tentorium base angle (TBA) was calculated between the sagittal tentorial orientation (vermis [no. 22] to the internal occipital protuberance [no. 33]) and the posterior cranial base (sella to basion). As a measure of the coronal tentorial orientation, the coronal tentorial angle (CTA) was calculated between the left and right line segments, passing through the most superior point of the cerebellar vermis (no. 22) and most lateral left and right cerebellar points (nos. 24 and 25, respectively). Furthermore, we calculated the cephalic index as the maximum brain width divided by the maximum brain length (Table 3), as an index of the general endocast morphology. The maximum width and maximum length were measured on aligned T1-weighted images in the coronal and the sagittal plane, respectively.

### Measurement error

Intraobserver error was calculated for measurements taken twice in the brain extraction step for 18 specimens, using one-way analysis of variance (ANOVA) for univariate measurements and the Procrustes ANOVA for Procrustes superimposed landmark coordinates. Procrustes ANOVA is analogous to two-way ANOVA (Klingenberg & McIntyre, 1998; Klingenberg et al., 2002). In this method, the deviation of the Procrustes superimposed landmark coordinates from the mean shape is divided into individual and asymmetric shape variation (objective asymmetry), the interaction of the individual symmetric variation and the asymmetric variation (fluctuating asymmetry), and the residual component (measurement error) (Klingenberg & McIntyre, 1998; Klingenberg et al., 2002).

### Data analysis

The symmetric variation in the Procrustes coordinates of all landmarks, i.e., those on the endocast surface, in the cranial base, and inside the brain structure, was analyzed by PCA. Principal component (PC) loadings were visualized using arrow vectors, which were set as the landmark mean ± 3 standard deviations. Shape variation along the PCs was illustrated by a combination of arrows and line segments, by connecting the original landmarks with a solid line (arrow tail) and transformed landmarks with a dashed line (arrow tip); this was carried out for the corpus callosum (nos. 1 and 2), brain stem (nos. 3–7), cerebellar vermis (nos. 8, 21, 22, and 23), lateral ventricles (nos. 9–14), midsagittal cranial base (nos. 28– 30), petrosal bone (nos. 26, 27, and 34–37), and sagittal (nos. 22 and 33) and coronal (nos. 22, 24, and 25) orientations of the tentorium cerebelli (Table 3). All the arrows and lines were colorized according to region: the cranial base (region A, blue), the surface of the infratentorial region (B, red), the inside of the infratentorial region (B, orange), the surface of the supratentorial region (C, dark green), and the inside of the supratentorial region (C, green) (Figure 2).

Ontogenetic trajectories along the first, second, and third PCs were visualized in two-dimensional plots. The ontogenetic trajectories of the angles, the relative sizes, and the cephalic index were compared between age stages. Differences between age stages were confirmed by Tukey–Kramer tests. The relationships between the relative sizes and shape variation were assessed by calculating correlation coefficients. The correlations of the observed shape changes and the relative sizes reflect the size–shape relationships during ontogeny. Furthermore, to visualize the ontogenetic changes in the cranial angles along the PC vectors, the angles of the petrosal bone and the tentorium cerebelli (the IPA, TBA, and CTA) were plotted by age stage, where the angles were measured in each age stage along the respective PCs. Along PC1; for example, the average angles in age stages 1–6 were calculated from the coordinates corresponding to the mean PC1 scores of each age stage, with the other PCs having grand mean values that were fixed at 0. The angles along PC2 and PC3 were calculated and visualized in the same manner.

## Results

### Measurement error

The results of a one-way ANOVA of the univariate measurements indicated that the variance in the repeated measurements for one specimen was significantly lower than the interindividual variation in all the univariate measurements (*p* < 0.01). The measurement error in the Procrustes superimposed landmark coordinates was also negligible and significantly smaller than for the other components according to the Procrustes ANOVA (Table 4).

**Table 4.** Results of one-way ANOVA of the “whole centroid size” and Procrustes ANOVA of shape, performed to assess measurement error. “Individual” refers to the symmetric shape variation. “Side” reflects directional asymmetry. “Individual * Side” reflects fluctuating asymmetry. SS: sum of squares, MS: mean squares, df: degree of freedom, F: F statistic, *p*: *p*-value.

| Effect | SS | MS | df | F | $p$ |
| --- | --- | --- | --- | --- | --- |
| Centroid size: |  |  |  |  |  |
| Individual | 437895.25 | 25758.544 | 17 | 5468.93 | <.0001 |
| Measurement error | 84.779679 | 4.709982 | 18 |  |  |
| Shape, Procrustes ANOVA: |  |  |  |  |  |
| Individual | 0.1036491 | 2.242E-05 | 4624 | 8.71 | <.0001 |
| Side | 0.0220356 | 9.068E-05 | 243 | 35.25 | <.0001 |
| Ind * Side | 0.010627 | 2.573E-06 | 4131 | 2.17 | <.0001 |
| Measurement error | 0.0109751 | 1.184E-06 | 9270 |  |  |

### Ontogenetic changes in overall size, relative sizes, and shape metrics

Figure 3 shows the ontogenetic trajectories of the whole CS of the landmarks, the relative sizes, the angle measurements, and the cephalic index. The whole CS increases rapidly over the first few years (stages 1–3), and increases gradually thereafter (Figure 3(a)). The IRE undergoes a moderate change during early postnatal ontogeny (stages 1– 3) and decreases thereafter (stages 3–6), indicating that expansion of the cranial base occurs late in ontogeny relative to the whole brain (Figure 3(b)). This is consistent with a previous observation (Zollikofer et al., 2017). The RIE increases shortly after birth (stages 1–3) and decreases gradually during stages 3–6 (Figure 3(c)). The IDE undergoes a dramatic increase, associated with expansion of the infratentorial region (stages 1–3), and then changes moderately (stages 3–6) (Figure 3(d)). This trend corresponds to the expansion of the cerebellum in the early stage of human postnatal development (Knickmeyer et al., 2008).

**Figure 3.**
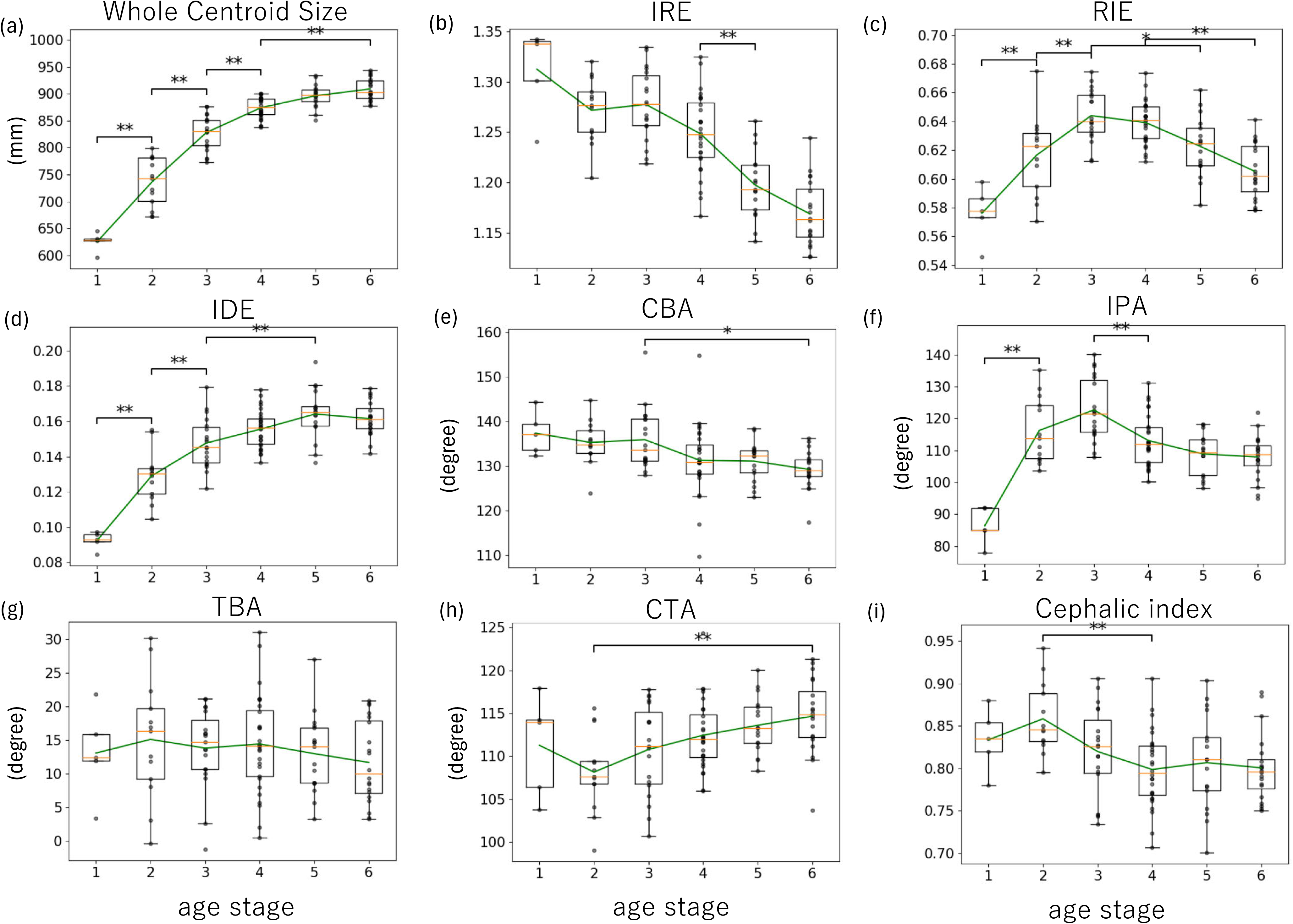
Ontogenetic changes in size and shape indices. (a) Centroid size of the landmarks; (b) index of relative encephalization (IRE); (c) index of relative infratentorial encephalization (RIE); (d) index of differential encephalization inside the brain (IDE); (e) cranial base angle (CBA); (f) inter-petrosal angle (IPA); (g) tentorium base angle (TBA); (h) coronal tentorial angle (CTA); (i) cephalic index. Green lines connect the mean values of the age stages. P-values of the differences between age stages according to the Tukey–Kramer test are indicated (*p < 0.05, **p < 0.01).

The CBA shows a moderate decrease in all age stages, and the difference in the CBA is significant only between stages 3 and 6 (*p* < 0.05) (Figure 3(e)). This result is different from the previous observation that the human cranial base flexes rapidly up to the postnatal age of 2 years and remains constant in late ontogeny (Lieberman & McCarthy, 1999). In this study, the sella point (no. 29) was located in the center of the pituitary gland (Table 3) and not in the center of the sella turcica, as in a previous study (Lieberman & McCarthy, 1999). Additionally, the most inferior point of the foramen caecum of the anterior cranial base was located on the basis of the observations of the soft tissues in this study, potentially differing from the point acquired on the bone surface in Lieberman and McCarthy (1999); this difference is possibly caused by the difference in the observations of the soft and hard tissue. The IPA increases dramatically during early ontogeny (stages 1–3), decreases slightly in stages 3–4, and remains constant thereafter (stages 4–6) (Figure 3(f)). This is consistent with a previous study showing that the human petrosal bone reorients coronally after birth (Dean & Wood, 1984). The TBA remains almost constant during growth (Figure 3(g)), and the CTA undergoes a moderate change in early ontogeny and slightly increases thereafter (Figure 3(h)) despite the size of the cerebellum under the tentorium cerebelli increasing rapidly in early ontogeny. The cephalic index decreases gradually during early development (stages 2–4) (Figure 3(i)), which means that the endocast becomes longer relative to its width.

Correlation coefficients of the two angle measurements (the CBA and IPA) with various indices (the whole CS, IRE, RIE, IDE, and cephalic index) were calculated. The CBA correlates positively with the IRE (*r* = 0.39, *p* < 0.001), and negatively with the whole CS (*r* = −0.35, *p* < 0.001) and the IDE (*r* = −0.37, *p* < 0.001). The IPA correlates positively with the RIE (*r* = 0.52, *p* < 0.001) (Table 5).

**Table 5.** Correlations between indices of shape (the CBA, IPA, and PCs 1–3) and size (the whole CS, IRE, IDE, and cephalic index). PC3 was divided into early and late stages because of its curvilinear trajectory. (\**p* < 0.01, \*\**p* < 0.001) CBA: cranial base angle, IPA: inter-petrosal angle, CS: centroid size, IRE: index of relative encephalization, RIE: index of relative infratentorial encephalization, IDE: index of differential encephalization inside the brain, PC: principal component.

|  | whole CS | IRE | RIE | IDE | Cephalic index |
| --- | --- | --- | --- | --- | --- |
| CBA | -0.352** | 0.394** | 0.0458 | -0.366* | 0.112 |
| IPA | 0.129 | 0.210 | 0.518** | 0.216 | 0.193 |
| PC1 | 0.827 ** | -0.650** | 0.157 | 0.808** | -0.555** |
| PC2 | -0.141 | 0.472** | 0.177 | -0.314* | -0.790** |
| PC3 | -0.188 | -0.335** | -0.678** | -0.239 | -0.0247 |
| PC3 (stage 1-3) | -0.829** | 0.0591 | -0.798** | -0.756** | 0.148 |
| PC3 (stage 3-6) | 0.477** | -0.651** | -0.627** | 0.213 | -0.0862 |

### Ontogenetic changes in the whole brain and neurocranium

The shape variation in the ontogenetic series is illustrated by the plots of the PC scores. Figure 3 shows plots of the first three PCs: PC1, PC2, and PC3 explain 32.0%, 18.7%, and 11.6% of the variance in shape, respectively, summing to 62.3%. Ontogenetic shape changes are clearly visible in PC1 and PC3. The patterns of the shape changes along the PCs are visualized in Figure 5. Shape changes along PCs are also shown in a movie (Video S1).

PC1 shows an almost linear developmental pattern throughout the ontogeny, where the PC1 scores are significantly different between age stages 2–5 (*p* < 0.01). PC1 strongly correlates with the whole CS (*r* = 0.83, *p* < 0.001), IRE (*r* = −0.65, *p* < 0.001), IDE (*r* = 0.81, *p* < 0.001) and cephalic index (*r* = −0.56, *p* < 0.001) (Table 5). The shape change between the lower and higher PC1 scores corresponds to the decrease in endocranial height and increase in length by projection of the frontal pole, the anterior and superior shift of the anterior floor of the cranial base, posterolateral elongation of the petrous bone, and relative widening and posterior expansion of the posterior cranial fossa. The brain stem position changes little with changes in the posterior cranial base in the sagittal view. By contrast, posterior changes are seen in the cerebellar vermis and tentorium cerebelli positions, and the lateral ventricles and corpus callosum orientations change in the sagittal view (Figure 5(a)). PC2 shows little ontogenetic change and seems to exhibit mostly individual variation, with a statistically significant difference only seen between age stages 1 and 5, and between stages 1 and 6 (*p* < 0.05) (Figure 4(a)). Shape changes between lower and higher PC2 scores are associated with increased endocranial length and decreased width; consistent with this result, PC2 shows a strong negative correlation with the cephalic index (*r* = −0.79, *p* < 0.01) (Figure 5(b), Table 5). Although the PC plots show that the endocast in age stage 1 is dolichocephalic relative to that in age stages 5 and 6, indicating a “brachiocephalic shift” during ontogeny (Figure 4(a)), the cephalic index decreases during ontogeny, and the endocranial length increases relative to its width (a “dolichocephalic shift”) (Figure 3(i)). This contradictory situation is potentially explained by the combination of shape changes along PC1 and 2. The dolichocephalic change during ontogeny is mainly reflected in PC1 (Figure 5(a), Table 5); concerning PC2, a brachiocephalic shift can be observed only in early ontogeny (Figure 4(a)). The shape change in brain structures along PC2 is minimal in the sagittal view. By contrast, the midsagittal anterior cranial base shows an inferior shift in position, and the inferior horn of the lateral ventricles shifts inwards in the coronal and transversal views (Figure 5(b)).

**Figure 4.**
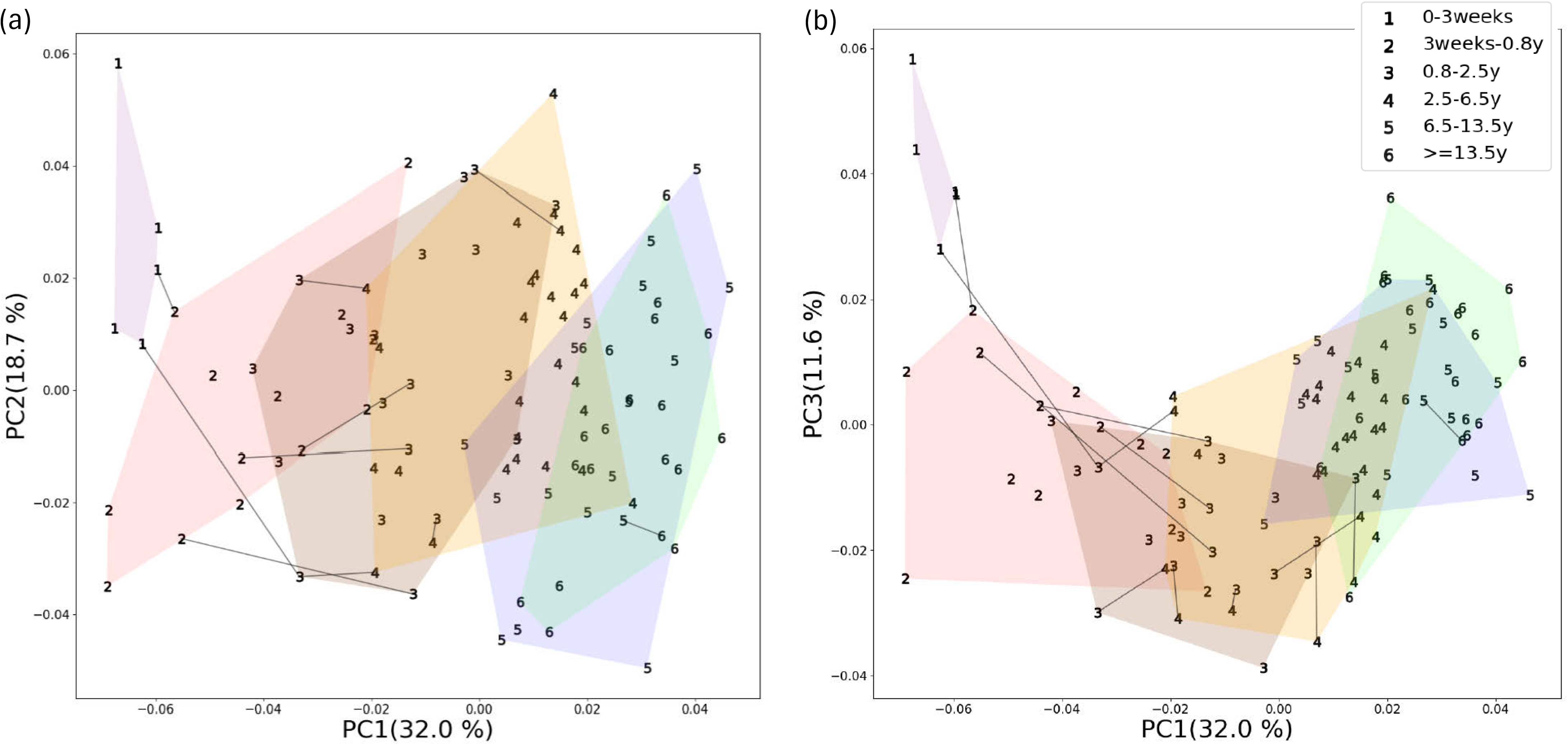
Plots of the first three principal components (PCs). (a) PC1 vs. PC2; (b) PC1 vs. PC3. Black lines connect the values of the different age stages for longitudinal specimens.

**Figure 5.**
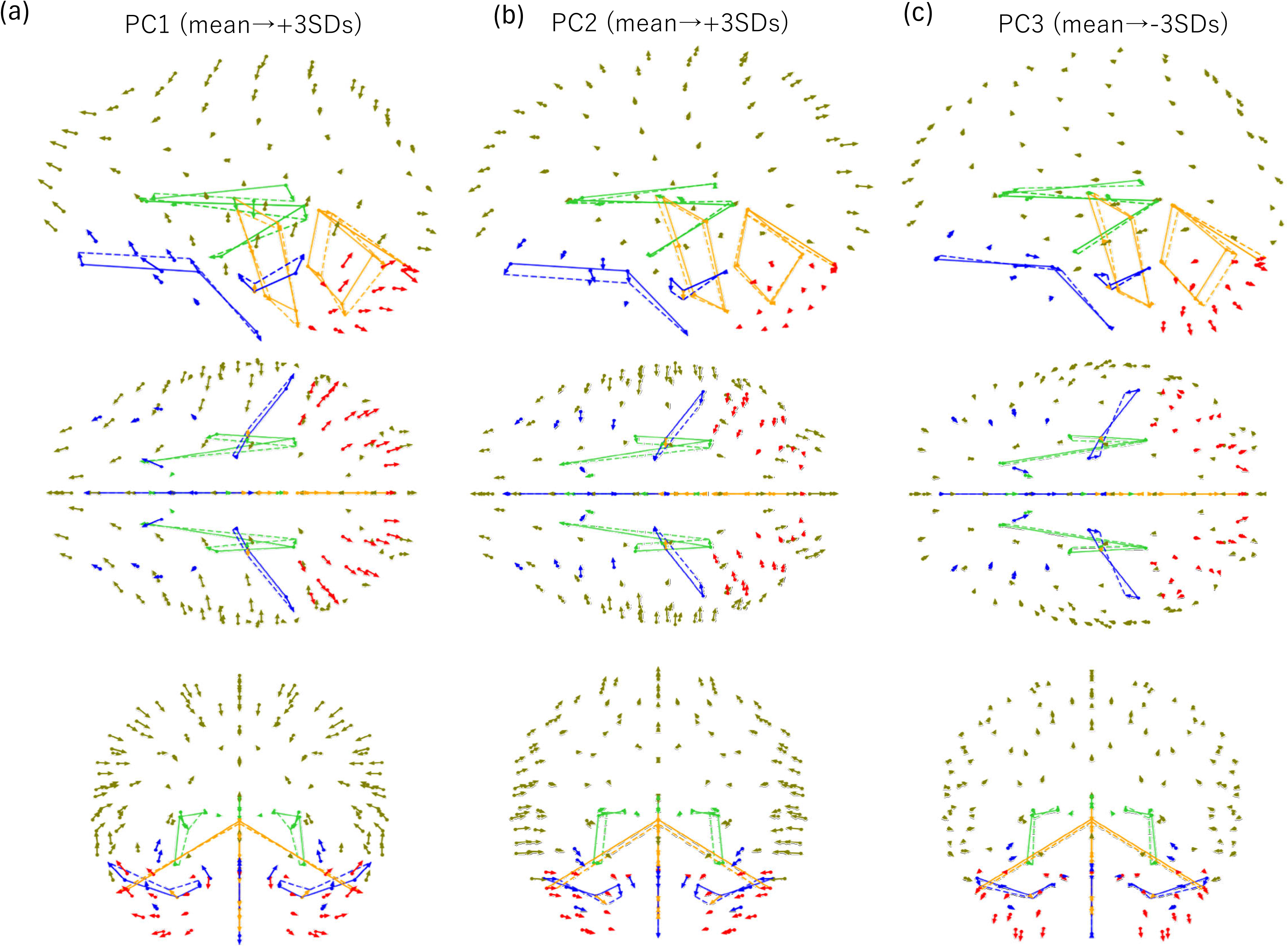
Shape variations along the principal components (PCs). (a) Shape changes along PC1, (b) PC2, and (c) PC3. Arrows connect the mean ± 3 standard deviation (SD) values. Landmark colors indicate the area: cranial base (region A, blue), surface of the infratentorial region (region B, red), inside of the infratentorial region (region B, orange), surface of the supratentorial region (region C, dark green), and inside of the supratentorial region (region C, green). The solid line segments indicate the mean shape of the structures, and the dashed line segments indicate the transformed shapes.

The PC3 scores exhibit a non-linear growth course, significantly decreasing between age stages 1 and 2 (*p* < 0.01) and increasing between stages 3 and 5 (*p* < 0.01) (Figure 4(b)). The shape change between higher and lower PC3 scores corresponds to inferior and posterior expansion of the posterior cranial fossa and the cerebellar vermis. In addition, the petrous orientation changes more coronally, and the inferior horns of the lateral ventricles are located in the interior part. The sagittal orientation of the tentorium cerebelli shows slight rotation posteriorly and superiorly (Figure 5(c)). The shape change along PC3 largely corresponds to the previously reported globularization phenomenon (Neubauer et al., 2010; Zollikofer et al., 2017). In terms of the relative sizes, PC3 shows a strong negative correlation with the RIE (*r* = −0.68, *p* < 0.001) as well as a negative correlation with the IRE (*r* = −0.34, *p* < 0.001) (Table 5). In addition to the overall PC3 scores, we calculated correlation coefficients for the early (stages 1–3) and late (stages 3– 6) stages because of the curvilinear ontogenetic trajectory of PC3. PC3 shows strong negative (stages 1–3) and positive (stages 3–6) correlations with the whole CS, a negative correlation with the IRE in stages 3–6 , a negative correlation with the IDE in stages 1–3, and negative correlations with the RIE in both stages 1–3 and stages 3–6.

Figure 6 shows the ontogenetic changes in the three angle measurements (the IPA, TBA, and CTA) along the PCs, where each “ontogenetic point” was calculated from the average shape in each age stage along the PC vector. Although the magnitude of the ontogenetic shift for the three angles is small, the changes were distinct among the PCs. Both the IPA and the TBA increase rapidly in early ontogeny along PC3, and decrease gradually from age stage 3 onward, whereas along PC1 and 2 they change little. By contrast, the CTA increases steadily along PC1, although it changes little along PC2 and PC3. The overall shape change patterns of the age stages along PC1–3 are shown in a movie (Video S2).

## Discussion

### Ontogenetic changes in the human brain and the braincase

To understand human-specific developmental patterns of the brain and the cranium, previous studies endeavored to obtain meaningful information from both the brain and the braincase. Although it has been argued that we should be aware of direct or indirect associations of endocranial imprints with the brain sulci (Alatorre Warren et al., 2019; Dumoncel et al., 2021), it is still important to consider size–shape interactions during development, which gave rise to the exceptionally large human brain. To shed light on the mechanisms underlying previously proposed hypotheses, we measured ontogenetic shape changes in both bony and internal soft tissue structures during human postnatal development according to changes in size (the absolute and relative size of the brain and brain subdivisions), PC vectors, angle measurements, and the cephalic index. Our results suggested that the shape changes in the human endocranium and structures inside the brain during development can be decomposed into three PCs, all of which generally accord with spatial packing constraints between the cranial base and the entire brain, evidenced by the significant correlations of the IRE with the PCs (Table 5). The three PCs also showed dissociable associations with brain growth and changes in the size of the supratentorial (i.e., cerebrum) and infratentorial (i.e., cerebellum) spaces, and the cranial base.

PC1 showed a linear trajectory during the course of ontogeny, where the PC scores were significantly different between age stages. The endocranial shape changes along the PC1 were dolichocephalic, showing a negative correlation with the cephalic index (*r* = - 0.56), a positive correlation with the IDE (*r* = 0.81; greater expansion of the cerebellum relative to the cerebrum), and a negative correlation with the IRE (*r* = −0.65), largely corresponding to a previously reported shape change pattern inferred to be related to the expansion of the face and the cranial base relative to the brain (Zollikofer et al., 2017). Our results also showed that the shape of internal brain structures such as the corpus callosum, lateral ventricle, and cerebellar vermis exhibited moderate shifts along PC1. Furthermore, it was noteworthy that the coronal angle between the tentorium cerebelli (the CTA) increased monotonically along PC1 (Figure 6(c)).

**Figure 6.**
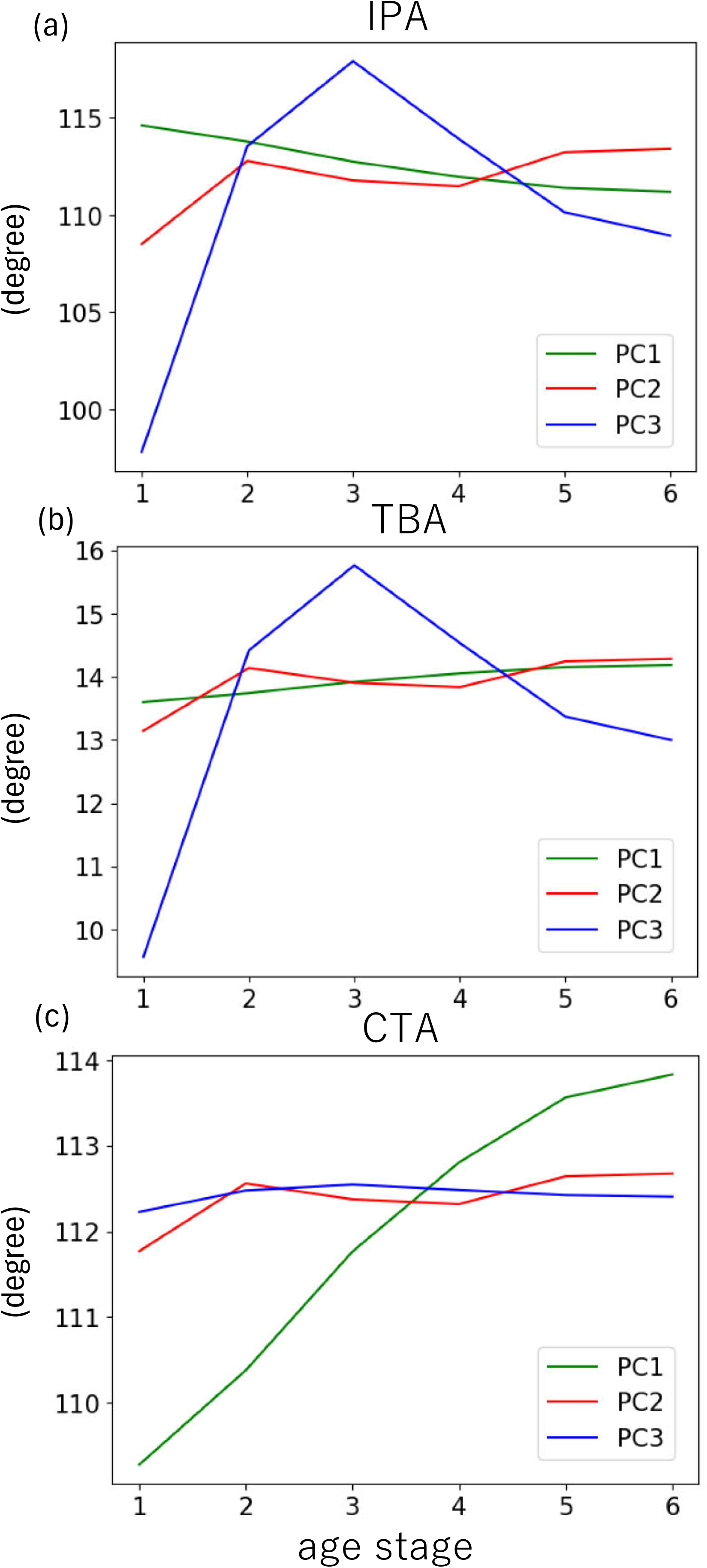
Angle sizes in each age stage along each principal component (PC) (green: PC1, red: PC2, blue: PC3). (a) The inter-petrosal angle (IPA), (b) tentorium base angle (TBA), and (c) coronal tentorial angle (CTA).

PC2 showed mostly idiosyncratic variation, reflecting ontogenetic changes only in the early period. The strong negative correlation with the cephalic index (*r* = −0.79) and the moderate positive correlation with the IRE (*r* = 0.47) indicate wide individual variation in brain shape, which can be partly explained by the general spatial packing hypothesis.

PC3 exhibited a unique curvilinear trajectory during postnatal development. Although the correlation with the whole CS was weak (*r* = −0.19), PC3 had stronger correlations with the CS values after dividing the data into earlier (stages 1–3, *r* = −0.83) and later stages (stages 3–6, *r* = 0.48). In stages 1–3, the PC3 scores decreased with age, whereas the infratentorial space (i.e., cerebellum) expanded greatly relative to the basicranium (the RIE) and the cerebrum (the IDE), which is consistent with a previous observation of relative expansion of the cerebellar volume in the first 2 years of life in humans (Knickmeyer et al., 2008). In stages 3–6, the PC3 scores in this study moderately increased with age, whereas the growth of the cerebrum and the cerebellum (i.e., the whole brain) slowed relative to the basicranial and facial components (as evidenced by the negative correlations with the IRE and RIE, but not with the IDE, in Table 5). Along the PC3 vector, the IPA and the TBA exhibited similar “up-and-down” trajectories, although they were largely stable along PC1 and PC2 (Figure 6(a) and 6(b)). These observations indicate that PC3 exhibits different ontogenetic orientations between the early and late stages of development, i.e., before and after stage 3, and the IRE and IDE only partly explain this shape change.

The ontogenetic shape changes reflected in the PC vectors were not fully compatible with the absolute IPA, TBA, and CTA values (Figure 3). The angles of the tentorium cerebelli remained almost constant during early postnatal ontogeny, including both the sagittal (the TBA; Figure 3(g)) and the coronal (the CTA; Figure 3(h)) angles, despite the dramatic expansion of the infratentorial region after birth (Figure 3(d)). These results are consistent with the previous observation that the sagittal orientation of the tentorium cerebelli and the occipital bone changes little postnatally in humans (Rehder et al., 2016). The inconsistency between the PC results and the absolute values can be explained by the absolute ranges of the angles and the use of PCA, which decomposes total variation into a few major independent components. For example, the absolute range of the IPA from stage 1 to 3 was approximately 35 degrees (Figure 3(f)), whereas that along PC3 was 25 degrees (Figure 6(a)), contributing 11% to the total variance explained for PC3. In other words, the shift in PC3 (25 degrees) can be considered large (> 70% of the absolute shift [35 degrees]) relative to the contribution to the variance explained. We can confirm that the large shift in the IPA angle from stage 1 to 3 along PC3 is characteristic of the shape change of the brain and the braincase in the early ontogenetic period seen with marked expansion of the cerebellum.

The absolute shifts, and those along the PC3 vector, of the TBA and CTA in this study are also interesting. Because these two angles reflect the sagittal and coronal orientations of the roof of the tentorium cerebelli, the different change patterns therein along the PC3 support developmental plasticity along the antero–posterior (or cranio–caudal) axis, and on the plane vertical to the axis. Although the absolute shifts in both the TBA and the CTA were small in early ontogeny (Figure 3(g, h)), the shift in the TBA (6 degrees) along PC3 was marked (Figure 6(b)), whereas that in the CTA was minimal along PC3 but increased continuously over the whole growth period along PC1 (Figure 6(c)). Therefore, we can assume that the developmental shape changes in early ontogeny along PC3 occur only along the axis of the cranium, and not in its coronal plane.

### Size–shape relationships in the developmental brain and the braincase

It has been argued that a human-specific ontogenetic change in the braincase is the relative expansion of the parietal and cerebellar regions seen in human neonates, described as globularization (Neubauer et al., 2010). However, Zollikofer et al. (2017) observed globularization not only in humans but also in gorillas and orangutans, albeit not in chimpanzees or bonobos. In addition, they found that the IRE (note their definition of the IRE is different to that in this study) can be considered to reflect “cranial integration” of intra- and inter-taxon variation in endocranial morphology, commonly being regarded as evidence of pervasive spatial packing constraints (Zollikofer et al., 2017).

In our human ontogenetic study, measures of relative size and shape, such as the CBA, IPA, and PC scores, showed interesting correlations (Table 5). The IRE values significantly correlated with all the three PC vectors, as well as with the CBA. We postulate that the IRE can serve as an effective measure of spatial packing constraints (Zollikofer et al., 2017), and ontogenetic shape variation in the human brain and braincase can also be explained by the spatial packing hypothesis. In addition, comparison of the correlations of the IRE with the PC3 scores, being almost zero in stages 1–3 versus the higher correlations in stages 3–6, indicate that spatial constraints are more influential in later stages.

The strong correlations of the RIE likely reflect the influence of infratentorial spatial packing constraints. The RIE showed significant correlations with the IPA, as well as with the PC3 scores during both the early and late stages (Table 5). From these results, we infer that the relative changes in size of the cerebellum and the cranial base underlie the shape changes along the PC3 vector, such as the reorientation of the petrosal bone and the inferior expansion of the posterior cranial fossa under the tentorium cerebelli with the moderate change in sagittal tentorial orientation. These results correspond to those of previous studies on the bony shape changes observed in humans during early ontogeny (Dean & Wood, 1984; Lieberman & McCarthy, 1999; Neubauer et al., 2010; Zollikofer et al., 2017).

### Study limitations

This study clarified the postnatal development of the human braincase and soft tissue structures and inferred developmental mechanisms on the basis of the relationships between measures of shape and relative size. The observed shape changes largely corresponded to those described as human-specific traits in previous reports (Dean & Wood, 1984; Lieberman & McCarthy, 1999; Neubauer et al., 2010; Zollikofer et al., 2017). We argue that the infratentorial spatial packing constraint between the cranial base and the infratentorial region during early ontogeny causes inferior expansion of the posterior cranial fossa, as a human-specific form of globularization. However, we only studied human postnatal ontogeny and cannot make inferences regarding inter-taxon variability.

The infratentorial spatial packing constraint may differ between primate species. In chimpanzees and bonobos, postnatal globularization is not seen (Neubauer et al., 2010; Zollikofer et al., 2017), although these species have a larger cerebellum relative to the total brain volume than humans (Rilling & Insel, 1998; Semendeferi & Damasio, 2000). Relative to the postnatal period, the sagittal section of the human tentorium cerebelli rotates inferoposteriorly during the fetal period, and this rotation is correlated with enlargement of the supratentorial volume relative to the infratentorial volume (Moss et al., 1956; Jeffery, 2002; Matsunari et al., 2023). Therefore, the relative sizes of the cerebrum, the cerebellum, and the cranial base during the fetal period affect the tentorial position and orientation at birth, and the inter-taxon differences in prenatal development potentially impose spatial constraints on postnatal cerebellar expansion that are dissociable between primate species. Inter-taxon variation in other aspects of cerebellar organization, such as the relative size of the cerebellar vermis and the cerebellar hemispheres (MacLeod et al., 2003; MacLeod, 2012), may also be influential. Interspecific comparisons are needed to reveal the factors driving developmental variation across species, and to interpret this variation from an evolutionary perspective.

## Acknowledgements

The data and/or research tools used in the preparation of this manuscript were obtained from the NIMH Data Archive, a collaborative informatics system created by the National Institutes of Health as a national resource to support and accelerate research in mental health (dataset identifier: 10.15154/vwwz-mt68). This manuscript reflects the views of the authors and may not reflect the opinions or views of the NIMH or those submitting original data to the NIMH Data Archive. We thank Michael Irvine, PhD, from Edanz (https://jp.edanz.com/ac) for editing a draft of this manuscript.

## Author contributions

K.T. designed the research, performed the analyses, and wrote the initial draft. O.K. contributed to the analysis and wrote the manuscript.

---

**Video S1.**
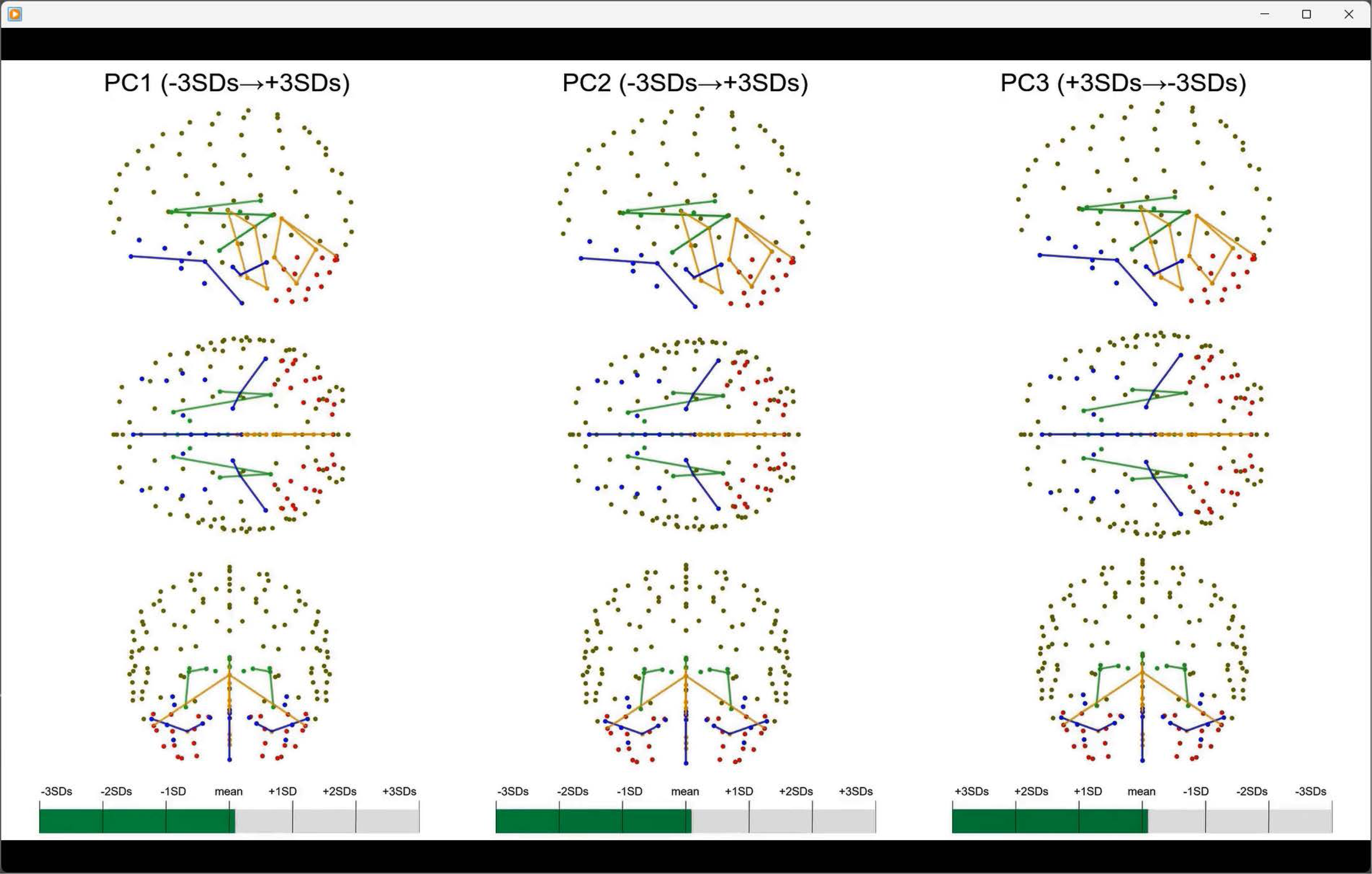
Shape changes along principal components (PCs).

**Video S2.**
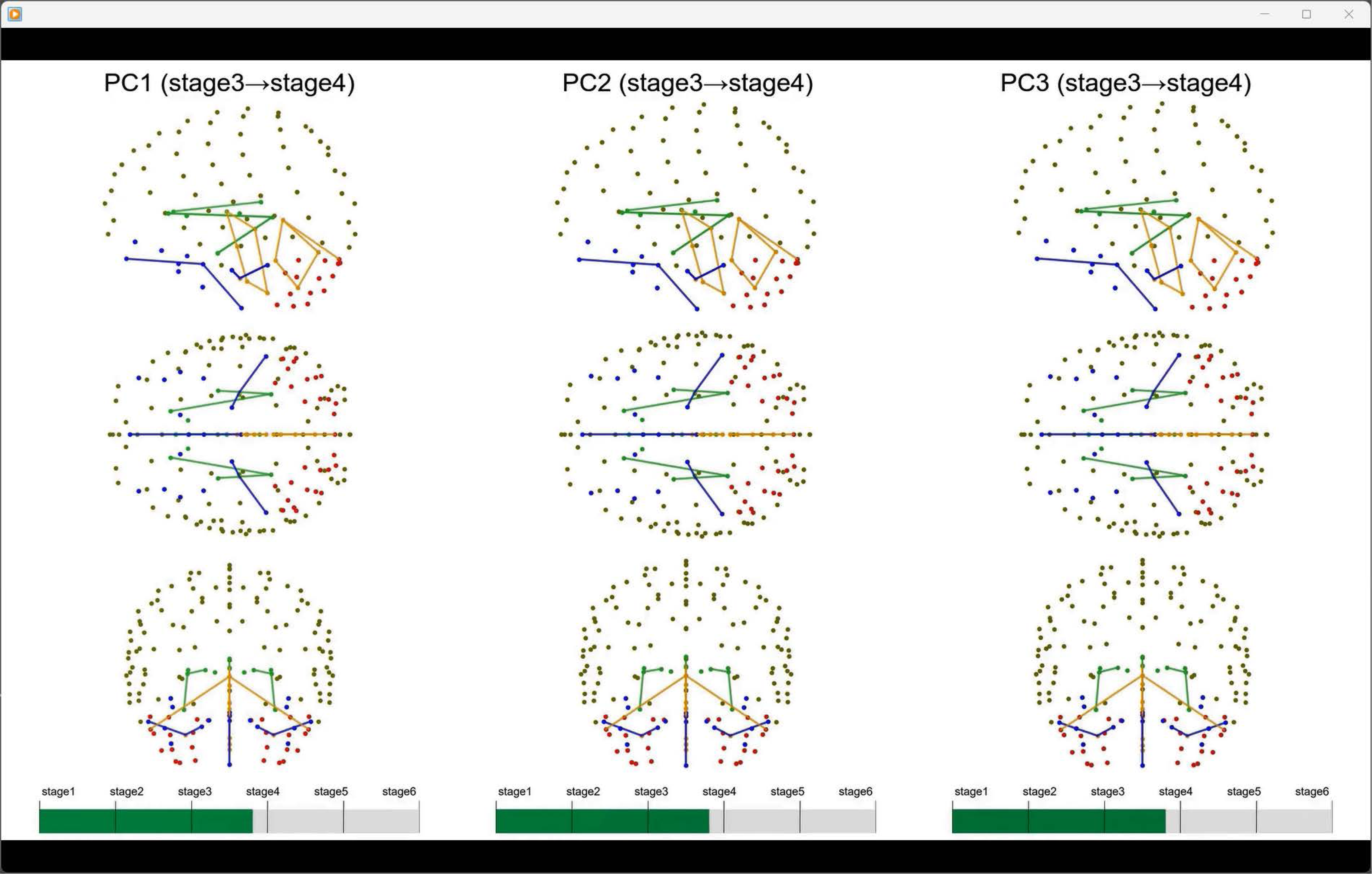
Shape changes along age stages 1–6.

## Notes

### Competing Interest Statement

The authors have declared no competing interest.

## References

Alatorre Warren JL, Ponce de León MS, Hopkins WD, Zollikofer CP (2019) Evidence for independent brain and neurocranial reorganization during hominin evolution. Proceedings of the National Academy of Sciences 116, 22115–22121.

Aldridge K (2011) Patterns of differences in brain morphology in humans as compared to extant apes. Journal of Human Evolution 60, 94–105.

AlQahtani SJ, Hector MP, Liversidge HM (2010) Brief communication: The London atlas of human tooth development and eruption. American Journal of Physical Anthropology 142, 481–490.

Ashburner J, Friston KJ (2005) Unified segmentation. NeuroImage 26, 839–851.

Barkovich AJ, Kjos BO, Jackson Jr DE, Norman D (1988) Normal maturation of the neonatal and infant brain: MR imaging at 1.5 T. Radiology 166, 173–180.

Barton RA, Harvey PH (2000) Mosaic evolution of brain structure in mammals. Nature 405, 1055–1058.

Barton RA, Venditti C (2013) Human frontal lobes are not relatively large. Proceedings of the National Academy of Sciences 110, 9001–9006.

Bastir M, Rosas A, Gunz P, et al. (2011) Evolution of the base of the brain in highly encephalized human species. Nature Communications 2, 588.

Bird CR, Hedberg M, Drayer BP, Keller PJ, Flom RA, Hodak JA (1989) MR assessment of myelination in infants and children: usefulness of marker sites. American Journal of Neuroradiology 10, 731–740.

Botton-Divet L, Cornette R, Fabre A, Herrel A, Houssaye A (2016) Morphological analysis of long bones in semi-aquatic mustelids and their terrestrial relatives. Integrative and Comparative Biology 56, 1298–1309.

Bruner E, Amano H, de la Cuétara JM, Ogihara N (2015) The brain and the braincase: a spatial analysis on the midsagittal profile in adult humans. Journal of Anatomy 227, 268–276.

Bruner E, Manzi G, Arsuaga JL (2003) Encephalization and allometric trajectories in the genus Homo: evidence from the Neandertal and modern lineages. Proceedings of the National Academy of Sciences 100, 15335–15340.

Bruner E, Preuss TM, Chen X, Rilling JK (2017) Evidence for expansion of the precuneus in human evolution. Brain Structure and Function 222, 1053–1060.

Bull JW (1969) Tentorium cerebelli. Proc R Soc Med 62, 1301–1310.

Dean MC, Wood BA (1984) Phylogeny, neoteny and growth of the cranial base in hominoids. Folia Primatologica 43, 157–180.

Dean MC (1988) Growth processes in the cranial base of hominoids and their bearing on morphological similarities that exist in the cranial base of Homo and Paranthropus. In: Evolutionary History of the ‘Robust’ Australopithecines. (ed. Grine FE), pp.107– 112, New York: Aldine de Gruyter.

Despotović I, Goossens B, Philips W (2015) MRI segmentation of the human brain: challenges, methods, and applications. Computational and Mathematical Methods in Medicine 2015.

Dumoncel J, Subsol G, Durrleman S, et al. (2021) Are endocasts reliable proxies for brains? A 3D quantitative comparison of the extant human brain and endocast. Journal of Anatomy 238, 480–488.

Evans AC, Brain Development Cooperative Group (2006) The NIH MRI study of normal brain development. NeuroImage 30, 184–202.

Falk D (1985) Hadar AL 162-28 endocast as evidence that brain enlargement preceded cortical reorganization in hominid evolution. Nature 313, 45–47.

Friede H (1981) Normal development and growth of the human neurocranium and cranial base. Scandinavian Journal of Plastic and Reconstructive Surgery 15, 163–169.

Godfrey L, Jacobs KH (1981) Gradual, autocatalytic and punctuational models of hominid brain evolution: A cautionary tale. Journal of Human Evolution 10, 255–272.

Gómez-Robles A, Hopkins WD, Sherwood CC (2014) Modular structure facilitates mosaic evolution of the brain in chimpanzees and humans. Nature Communications 5, 4469.

Gould SJ (1977) Ontogeny and phylogeny. Cambridge, MA: Belknap Press of Harvard University Press.

Gunz P, Mitteroecker P, Bookstein FL (2005) Semilandmarks in three dimensions. In: Modern morphometrics in physical anthropology. (ed. Slice DE), pp.73–98, New York: Kluwer Academic/Plenum Publishers.

Gunz P, Neubauer S, Golovanova L, Doronichev V, Maureille B, Hublin J (2012) A uniquely modern human pattern of endocranial development. Insights from a new cranial reconstruction of the Neandertal newborn from Mezmaiskaya. Journal of Human Evolution 62, 300–313.

Gunz P, Neubauer S, Maureille B, Hublin J (2010) Brain development after birth differs between Neanderthals and modern humans. Current Biology 20, R921–R922.

Herculano-Houzel S (2007) Encephalization, neuronal excess, and neuronal index in rodents. The Anatomical Record: Advances in Integrative Anatomy and Evolutionary Biology: Advances in Integrative Anatomy and Evolutionary Biology 290, 1280–1287.

Herculano-Houzel S (2012) The remarkable, yet not extraordinary, human brain as a scaled-up primate brain and its associated cost. Proceedings of the National Academy of Sciences 109, 10661–10668.

Herculano-Houzel S (2009) The human brain in numbers: a linearly scaled-up primate brain. Frontiers in Human Neuroscience 3, 857.

Holloway RL (1983) Cerebral brain endocast pattern of Australopithecus afarensis hominid. Nature 303, 420–422.

Jeffery N (2003) Brain expansion and comparative prenatal ontogeny of the non-hominoid primate cranial base. Journal of Human Evolution 45, 263–284.

Jeffery N (2002) Differential regional brain growth and rotation of the prenatal human tentorium cerebelli. Journal of Anatomy 200, 135–144.

Jeffery NS, Humphreys C, Manson A (2022) A human craniofacial life-course: cross-sectional morphological covariations during postnatal growth, adolescence, and aging. The Anatomical Record 305, 81–99.

Jeffery N, Spoor F (2002) Brain size and the human cranial base: a prenatal perspective. American Journal of Physical Anthropology: The Official Publication of the American Association of Physical Anthropologists 118, 324–340.

Jerison HJ (1977) The theory of encephalization. Annals of the New York Academy of Sciences 299, 146–160.

Jernigan TL, Brown TT, Hagler Jr DJ, et al. (2016) The pediatric imaging, neurocognition, and genetics (PING) data repository. NeuroImage 124, 1149–1154.

Klingenberg CP (2011) MorphoJ: an integrated software package for geometric morphometrics. Molecular Ecology Resources 11, 353–357.

Klingenberg CP, Barluenga M, Meyer A (2002) Shape analysis of symmetric structures: quantifying variation among individuals and asymmetry. Evolution 56, 1909–1920.

Klingenberg CP, McIntyre GS (1998) Geometric morphometrics of developmental instability: analyzing patterns of fluctuating asymmetry with Procrustes methods. Evolution 52, 1363–1375.

Knickmeyer RC, Gouttard S, Kang C, et al. (2008) A structural MRI study of human brain development from birth to 2 years. Journal of Neuroscience 28, 12176–12182.

Kochiyama T, Ogihara N, Tanabe HC, et al. (2018) Reconstructing the Neanderthal brain using computational anatomy. Scientific Reports 8, 6296.

Lieberman DE, McCarthy RC (1999) The ontogeny of cranial base angulation in humans and chimpanzees and its implications for reconstructing pharyngeal dimensions. Journal of Human Evolution 36, 487–517.

MacLeod C (2012) The missing link: evolution of the primate cerebellum. Progress in Brain Research 195, 165–187.

MacLeod CE, Zilles K, Schleicher A, Rilling JK, Gibson KR (2003) Expansion of the neocerebellum in Hominoidea. Journal of Human Evolution 44, 401–429.

Matsunari C, Kanahashi T, Otani H, et al. (2023) Tentorium cerebelli formation during human embryonic and early fetal development. The Anatomical Record 306, 515–526.

Moss ML (1958) The pathogenesis of artificial cranial deformation. American Journal of Physical Anthropology 16, 269–286.

Moss ML, Noback CR, Robertson GG (1956) Growth of certain human fetal cranial bones. American Journal of Anatomy 98, 191–204.

Moss ML, Young RW (1960) A functional approach to craniology. American Journal of Physical Anthropology 18, 281–292.

Neubauer S, Gunz P, Hublin J (2009) The pattern of endocranial ontogenetic shape changes in humans. Journal of Anatomy 215, 240–255.

Neubauer S, Gunz P, Hublin J (2010) Endocranial shape changes during growth in chimpanzees and humans: a morphometric analysis of unique and shared aspects. Journal of Human Evolution 59, 555–566.

Passingham RE, Smaers JB (2014) Is the prefrontal cortex especially enlarged in the human brain? Allometric relations and remapping factors. Brain, Behavior and Evolution 84, 156–166.

Ponce de León MS, Bienvenu T, Akazawa T, Zollikofer CP (2016) Brain development is similar in Neanderthals and modern humans. Current Biology 26, R665–R666.

Rehder R, Yang E, Cohen AR (2016) Variation of the slope of the tentorium during childhood. Child’s Nervous System 32, 441–450.

Rilling JK (2006) Human and nonhuman primate brains: are they allometrically scaled versions of the same design?*. Evolutionary Anthropology: Issues, News, and Reviews: Issues*, News, and Reviews 15, 65–77.

Rilling JK, Glasser MF, Preuss TM, et al. (2008) The evolution of the arcuate fasciculus revealed with comparative DTI. Nature Neuroscience 11, 426–428.

Rilling JK, Insel TR (1999) The primate neocortex in comparative perspective using magnetic resonance imaging. Journal of Human Evolution 37, 191–223.

Rilling JK, Insel TR (1998) Evolution of the cerebellum in primates: differences in relative volume among monkeys, apes and humans. Brain Behavior and Evolution 52, 308–314.

Ross CF, Ravosa MJ (1993) Basicranial flexion, relative brain size, and facial kyphosis in nonhuman primates. American Journal of Physical Anthropology 91, 305–324.

Ruff CB, Trinkaus E, Holliday TW (1997) Body mass and encephalization in Pleistocene Homo. Nature 387, 173–176.

Sandler HC (1944) The eruption of the deciduous teeth. The Journal of Pediatrics 25, 140–147.

Schlager S (2017) Morpho and Rvcg–shape analysis in R: R-packages for geometric morphometrics, shape analysis and surface manipulations. In: Statistical shape and deformation analysis. (eds. Zheng G, Li S, Székely G), pp.217–256 Academic Press.

Schoenemann PT, Sheehan MJ, Glotzer LD (2005) Prefrontal white matter volume is disproportionately larger in humans than in other primates. Nature Neuroscience 8, 242–252.

Scott JA, Grayson D, Fletcher E, et al. (2016) Longitudinal analysis of the developing rhesus monkey brain using magnetic resonance imaging: birth to adulthood. Brain Structure and Function 221, 2847–2871.

Semendeferi K, Damasio H (2000) The brain and its main anatomical subdivisions in living hominoids using magnetic resonance imaging. Journal of Human Evolution 38, 317–332.

Semendeferi K, Lu A, Schenker N, Damásio H (2002) Humans and great apes share a large frontal cortex. Nature Neuroscience 5, 272–276.

Smith SM (2002) Fast robust automated brain extraction. Human Brain Mapping 17, 143–155.

Standring S, Ellis, H. J., Healy, et al. (2005) Gray’s anatomy: the anatomical basis of clinical practice. 39th ed. Elsevier.

Weaver AH (2005) Reciprocal evolution of the cerebellum and neocortex in fossil humans. Proceedings of the National Academy of Sciences 102, 3576–3580.

White DD (2005) Size and shape of the cerebellum in catarrhine primates and plio-pleistocene fossil hominins: A paleoneurological analysis of endocranial casts. Dissertation, State University of New York at Albany.

Yushkevich PA, Piven J, Hazlett HC, et al. (2006) User-guided 3D active contour segmentation of anatomical structures: significantly improved efficiency and reliability. NeuroImage 31, 1116–1128.

Zollikofer CP, Bienvenu T, Beyene Y, et al. (2022) Endocranial ontogeny and evolution in early Homo sapiens: The evidence from Herto, Ethiopia. Proceedings of the National Academy of Sciences 119, e2123553119.

Zollikofer CP, Bienvenu T, Ponce de León MS (2017) Effects of cranial integration on hominid endocranial shape. Journal of Anatomy 230, 85–105.

